# Small Accelerations of the cell generate sufficient nuclear motion to modulate transcriptional activity, driving cellular response independent of matrix strain

**DOI:** 10.1101/2025.04.07.647583

**Authors:** Nina Nikitina, Jonathan Wadsworth, Matthew Goelzer, Madison Goldfeldt, Nurbanu Bursa, Sean Howard, Chase Crandall, Amevi Semodji, Anamaria G. Zavala, Stefan Judex, Janet Rubin, Trevor J. Lujan, Clare K Fitzpatrick, Clinton T Rubin, Aykut Satici, Gunes Uzer

## Abstract

The cell’s mechanical environment is a fundamental determinant of its activity. Ostensibly, the cellular response is dependent on interactions between extracellular matrix deformations and the cell adhesome. Low-intensity vibration (LIV) induces sinusoidal mechanical accelerations that stimulate mesenchymal stem cell (MSC) anabolism despite generating minimal matrix strain. In this study, we tested the hypothesis that accelerations of less than 1g cause nuclear motions relative to the cell membrane in adherent cells, resulting in elevated stresses in the cytoskeleton that promote transcriptional activity. Coupling a piezoelectric vibration platform with real-time microscopy, we applied a 0.7g, 90Hz LIV signal that oscillates the cell with displacements of up to ±11 µm. Live-cell tracking revealed that the sinusoidal vibrations caused the nucleus to move ±1.27 µm (17% of total displacement) out of phase with the cell membrane. Disruption of the LINC complex, which mechanically couples the nucleoskeleton to the cytoskeleton, doubled the magnitude of this relative motion, indicating that the nucleo-cytoskeletal configuration plays a major role in regulating nuclear motion. Consistent with a previously reported increase in nuclear stiffness caused by LIV, machine-learning-based image segmentation of confocal micrographs showed that LIV increased both apical and basal F-actin fiber numbers, generating a denser, more branched actin network near the nucleus. Following six 20 min bouts of LIV applied to MSC, RNA sequencing identified 372 differentially expressed genes. Upregulated gene sets were linked to F-actin assembly and focal adhesion pathways. Finite element simulations showed that nuclear stresses increased by LIV up to 18% were associated with nuclei flattening and a 30-50% increase in actin-generated forces. These findings demonstrate that low-intensity accelerations, independent of matrix strain, can directly activate a response of the nucleus, leading to cytoskeletal reorganization and heightened nuclear stresses. Thus, even very small oscillatory mechanical signals can markedly influence cell outcomes, establishing a mechanosensing pathway independent of extracellular strains.

## Introduction

Living organisms must continuously sense, respond, and adapt to their physical environments at cellular, tissue, and organ levels. Through organismal adaptation, mechanical signals are key for preserving physiological function in many systems including musculoskeletal,^1^ immune,^2^ cardiovascular, and cognition.^3^ These adaptations begin at the level of the cell, where specialized mechanosensory complexes convert exogenous mechanical signals into biologically relevant responses, triggering differential gene expression as well as adaptations in cytoskeletal^4^ and nucleoskeletal architecture.^5^

In pluripotent cells, mechanically triggered processes guide many functions, including fate selection, expansion and apoptosis.^6^ The loss of mechanical input has detrimental effects. For example, microgravity conditions shift stem cell lineage selection towards adipogenesis at the expense of osteoblastogenic and myogenic differentiation,^7^ resulting in weaker bones and increased fatty infiltration in muscles. To combat the effects of microgravity, astronauts have practiced rigorous resistive exercises that generate significant forces and cause large extracellular matrix strains yet these fail, on average, to deter deconditioning.^8^ These data suggest that cells may not only recognize high force and low frequency events associated with resistive exercise, but may also be influenced by other kind of mechanical factors.

For instance, bone, muscle, and other systems may benefit from even gentle exercise, such as walking or Tai Chi, highlighting that a cell’s ability to recognize and respond to mechanical events is not directly dependent on high extracellular stresses. Indeed, beyond the impact spikes and resultant forces generated by locomotion, the weight-bearing skeleton is bombarded by low-magnitude, high-frequency mechanical signals^9^ due to the high-frequency power spectrum of postural muscle contractions during even quiet standing, an activity that is not experienced in space.^10^ As many physiologic systems, like vestibular, hearing, and touch are designed to sense high-frequency signals, perhaps all cells have the capacity to respond to high frequency signals, facilitating a sensor system based on acceleration, rather than substrate deformations.

Low-Intensity Vibration (LIV) is a mechanical signal developed as a surrogate for the power spectrum of muscle contractability.^20^ Low impact (<1g) and high frequency (30-100Hz) characteristics of LIV make a viable strategy to deliver mechanical signals to people who have limited exercise ability, such as some with osteoporosis,^11^ cerebral palsy,^12^ or cancer-related bone lesions.^13^ In bone, LIV-induced bone matrix deformations are orders of magnitude below the osteogenic threshold set by models,^14^ yet are still anabolic to bone.^15^ In contrast to exercise which can induce bone deformations up to 3500 microstrain (µε),^16^ LIV induces strains in bone of less than 20µε.^17^ Not surprisingly, *in vivo* studies have demonstrated that bone response to LIV may be independent of strain generated in the bone matrix,^17,18^ implying that cells have the capacity to directly sense accelerations.

At the cellular level, LIV-induced dynamic accelerations increase cell proliferation,^19^ maintain differentiation potential under replicative aging,^20^ and restore mechanosignaling under simulated microgravity.^21^ These cell level effects are in part driven by LIV-induced increases in cell contractility. LIV, through Focal Adhesion Kinase (FAK) mediated RhoA activation,^22^ contribute to cell stiffening and osteogenic differentiation of MSCs.^23,24^ Pointing to a potential mechanism of how LIV-induced dynamic accelerations are sensed by cells, disrupting nucleo-cytoplasmic connectivity mediated by Linker of Nucleoskeleton and Cytoskeleton (LINC) prevents LIV-induced RhoA-activation,^22^ and fails to regulate cellular stiffness in both MSCs^25^ and breast cancer cells.^26^ In contrast, when cells are subjected to extracellular matrix strain, decoupling the nucleus from the cytoskeleton does not affect FAK activation or cell contractility,^27^ underscoring that nuclear mechanosensitivity plays a role in LIV-induced signaling, but not in strain-based mechanotransduction.

These data led us to hypothesize that rather than engaging in an *outside-in* mechanotransduction paradigm, LIV signaling is independent of matrix deformations and is instead mediated by the nuclear sensing of dynamic accelerations through an *inside-out* signaling.

To this end, we developed a piezo-electric LIV-platform integrated with a microscope system that allowed us to measure nuclear and cellular motions, in the presence and absence of LINC function, to investigate whether they alter nuclei-associated F-actin architecture and increase the expression of actin and focal adhesion-related genes. We also tested whether statistically derived generalized FE models can predict the magnitude of LIV-induced mechanical stresses within cell nuclei, an important step towards elucidating the mechanical mechanism by which vibrations drive cellular adaptations.

## Methods and Data Collection

### Isolation of MSCs and cell culture

Bone marrow derived MSCs (mdMSC) from 8-10 wk male C57BL/6 mice were isolated from multiple mouse donors and pooled, providing a heterogenous MSCs cell line. Briefly, tibial and femoral marrow were collected in RPMI-1640, 9% FBS, 9% HS, 100 μg/ml pen/strep and 12μM L-glutamine. After 24 hours, non-adherent cells were removed by washing with phosphate-buffered saline and adherent cells were cultured for 4 weeks. Passage 1 cells were collected after incubation with 0.25% trypsin/1 mM EDTA × 2 minutes and re-plated in a single 175-cm^2^ flask. After 1-2 weeks, passage 2 cells were re-plated at 50 cells/cm^2^ in an expansion medium (Iscove’s modified Dulbecco’s Medium (IMDM), 9% FBS, 9% HS, antibiotics, L-glutamine).

mdMSCs were re-plated every 1-2 weeks for two consecutive passages up to passage 5 and tested for osteogenic and adipogenic potential, and subsequently frozen. Calf serum (CS) was obtained from Atlanta Biologicals (Atlanta, GA). MSCs were maintained in IMDM with FBS (10%, v/v) and penicillin/streptomycin (100μg/ml).

These isolated MSC stocks were stably transduced with a doxycycline-inducible plasmid expressing a mCherry tagged dominant-negative KASH (dnKASH) domain.^21^ The dnKASH plasmid was lentiviral packaged as a generous gift from Dr. Daniel Conway (Addgene # 125554). For doxycycline-inducible dnKASH, lentivius supernatant was added to growth media with polybrene (5 μg/ml). Lentivirus growth media mixture was added to 50-70% confluent MSCs. Lentivirus media was replaced 48 hours later with selection media containing G418 (1mg/ml) for 5 days to select stably infected dnKASH MSCs, these MSCs were subsequently frozen. Calf serum (CS) was obtained from Atlanta Biologicals (Atlanta, GA). MSCs were maintained in IMDM with FBS (10%, v/v) and penicillin/streptomycin (100μg/ml). To disable LINC function, dnKASH cells were given growth media containing doxycycline ( Dox, 1 μg/ml) for 48 hours to induce mCherry-KASH and prevent actin linking to Nesprins on the nuclear envelope.^28^ Controls were not exposed to Dox.

### Quantification of nuclear motions

We developed a LIV bioreactor based on a P-841.3 piezoelectric actuator and E-610 controller (Physik Instrumente) (**Fig1.a**). The bioreactor, capable of modulating platform frequency and acceleration between 0–300 Hz and 0–1g, was integrated with a Zeiss LSM 900 confocal microscope equipped with an environmental chamber. Shown in **Fig.1b**, acceleration was measured with both an accelerometer (LSM6DSOX+LIS3MDL, Adafruit) and fluorescent particle tracking (F8824, Fisher Scientific). For fluorescent particle tracking (**Fig.1c**), total bead travel was captured by setting the camera exposure equal to the wavelength of the LIV input signal (∼11 milisecond, ms) and measured via ImageJ. To measure nuclear motions, MSCs were plated at 3,000 cells/cm^2^ in growth media. 24 hours later, cells were incubated with NucBlue Hoechst 33342 stain (Fisher Scientific) according to the manufacturer’s protocol to label nuclei. Similarly, cell membrane motions were measured using calcein AM (Invitrogen, 100nM). Using an ORCA-Flash4.0 digital CMOS Camera, the imaging area was set to 200x200 pixels with a 1 ms exposure time, allowing us to capture nuclear motions at 1,000 frames per second (fps). Prior to LIV, X-Y locations for 10 to 30 cells were labeled and stored. Immediately upon commencing LIV, each cell location was successively recorded at 1000 fps for 50 ms using Plan-Apochromat 10x/0.45 M27 lens.

**Figure 1:**
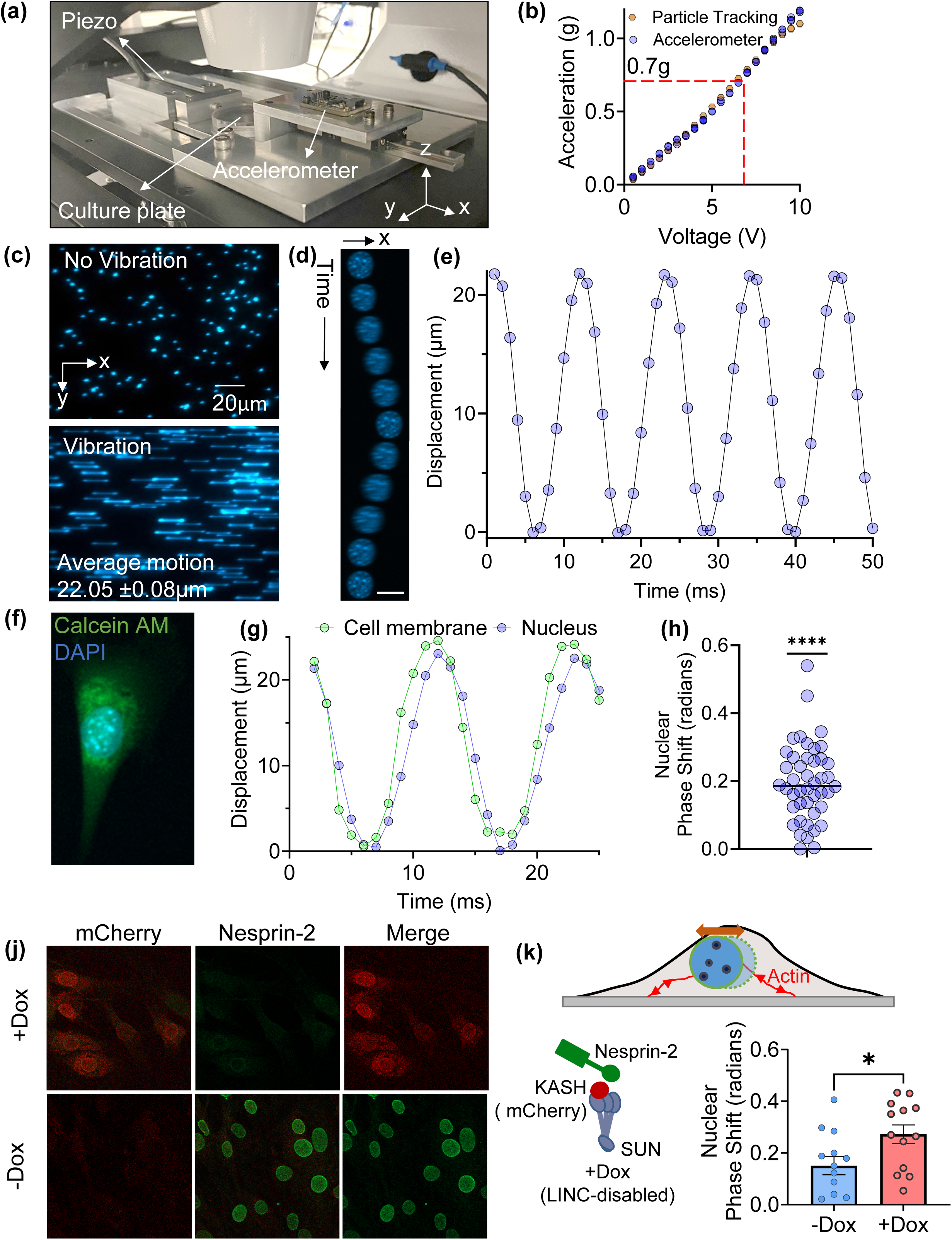
LIV generates an out of phase nuclear motion in a LINC complex dependent manner. **(a)** We developed an LIV bioreactor that can oscillate cells in a microscope chamber allowing us to horizontally vibrate a single 35mm imaging plate under microscopy with a **(b)** linear acceleration-voltage relationship with 0.7g, 90Hz LIV signal corresponding to approximately 6.5V (dashed red line). **(c)** Acceleration readings from an accelerometer were verified by tracking the time averaged travel path of fluorescent particles by setting the camera exposure equal to the wavelength of the input signal while **(d)** lowering the exposure time to 1ms allowed us to track the motion of a single nucleus vibrating at 90Hz, 0.7g over 11 frames (**e)** showing a periodic motion as expected. **(f)** Labeling the cell body with Calcein AM allowed us to **(g)** track the nucleus and the cell edge simultaneously, **(h)** revealing a nuclear phase shift of 0.2 ± 0.11 radians (∼0.35 ms average delay, N=47) corresponding to an averaged relative motion of 1.27µm during a 90Hz, 0.7g LIV. A one-sample Wilcoxon signed-rank test (GraphPad Prism) confirmed the lag was significantly different from zero (p < 0.0001). **(j)** Disabling LINC complex function during 90Hz, 0.7g vibrations in cells treated with +Dox to express mCherry-KASH **(k)** doubled the nuclear phase lag under LIV. An unpaired two-tailed Student’s t-test (GraphPad Prism) confirmed the increase was significant (**p < 0.05**).

Following the recording session, nuclear and membrane motions were tracked using the TrackMate plugin for ImageJ. The motion of the cell membrane was tracked automatically using the thresholding detector and the motion of the nucleus was tracked through the manual annotation setting with the location of the nucleus being found manually in each frame. A best fit sine wave was then generated for the membrane and nucleus and the phase shift between the two was determined and plotted (**Fig.1h**).

### Application of LIV regimen

To accommodate more than single culture plate, samples that did not require live imaging were subjected to LIV via a custom table-top vibration platform equipped with a Labworks ET-126 shaker. LIV was applied in 6 rounds. In each round, plates were placed on a machine at room temperature (RT) that vibrated them at 90 Hz, 0.7 g for 20 minutes. Control plates were not vibrated but were kept at RT while the other plates were being vibrated. Plates were incubated for 60 minutes at 37 °C between each round. Samples were taken 3 hours following the final round of LIV.

### RNA-seq

Total RNA was extracted using RNAeasy (Qiagen) for three samples per group. Total RNA samples were sent to Novogene for mRNA sequencing and analysis. Briefly, the index of the reference genome was built using Hisat2 v2.0.5 and paired-end clean 2 reads were aligned to the reference genome using Hisat2 v2.0.5. FeatureCounts v1.5.0-p3 was used to count the reads numbers mapped to each gene. Differential expression analysis was performed using the DESeq2 R package (1.20.0). DESeq2 provides statistical routines for determining differential expressions in digital gene expression data using a model based on the negative binomial distribution. The resulting P-values were adjusted using the Benjamini and Hochberg’s approach for controlling the false discovery rate. Genes with an adjusted p-value < 0.05 and fold-change (FC) > 0.2 found by DESeq2 were assigned as differentially expressed. Genes with significant differential gene expression were further analyzed with DAVID for pathway analysis^22^. Pathways with a p < 0.05 were selected.

### Immunofluorescence

Three hours after the last vibration, cells were fixed with 4% paraformaldehyde. Cells were permeabilized by incubation with 0.1% Triton X-100 for 5 minutes at 37°C. Staining was performed by incubating cells for 60 minutes at RT in a solution containing PBS with 1% Bovine Serum Albumin (blocking), Phalloidin (iFluor 488, Cayman Chemicals) for F-actin staining, and NucBlue Hoechst 33342 (Fisher Scientific) for nuclear staining. All staining was performed in a single step, protected from light to prevent photobleaching. Reagents used for immunofluorescence and their concentrations are listed in **Supplementary Table S1**. The fluorescent actin cytoskeleton images were obtained using a Zeiss LSM 900 microscope equipped with a Plan-Apochromat 40×/1.4 oil immersion objective and Zeiss Immersol 518 F immersion oil.

### Reconstruction of apical and basal actin stress fibers of msc from confocal microscope images

The reconstruction algorithm quantitatively characterizes the spatial organization of nuclei and associated actin stress fibers within the perinuclear area, our region of interest (ROI). The method routine consists of four phases: confocal image preprocessing, deep-learning-based segmentation, fiber and nuclear reconstruction, and visualization (Fig. S1).

#### Preprocessing (Fig. S1a)

Confocal images containing multiple cells were initially used to generate maximal intensity projections across all z-layers within the nuclear imaging channel. A U-Net deep learning model, previously trained, was applied to these projections to obtain masks outlining each nucleus, thereby locating individual perinuclear ROIs. Each nuclear outline served as a reference to extract both the nuclear region and the corresponding perinuclear actin fibers; each nucleus associated with its actin fiber was treated as a single instance. Nuclei touching image edges were excluded from the analysis.

The cropped image z-stacks of all individual nucleus–actin instances were rotated in the X–Y plane to align the predominant actin fiber orientation parallel to the X-axis. The rotation angle used was determined using a Hough Transform applied to maximal projections of an actin instance. Since apical and basal fibers in the perinuclear region of MSCs often differ in orientation, we reconstructed them separately by running the entire algorithm twice. For apical reconstruction, the algorithm pipeline used z-stacks rotated based on the apical fiber alignment. For basal reconstruction, the algorithm was re-run from the start using z-stacks rotated based on the basal fiber orientation. The apical fiber orientation was defined using the top half of the image layers, and the basal fiber orientation was defined using the bottom half. In the concluding step, the rotated sets of nuclei and F-actin individual instances were transformed into Y-Z cross-sectional 2D image layer sets along the X-axis.

#### Segmentation (Fig. S1b)

Actin fibers on the Y-Z cross-sectional images were visible as distinct dots in the image segmentation phase. These dots differed in size, shape, and intensity across the X-axis stacked layers, eliminating the possibility of using a single detection threshold for all. Adjusting the threshold manually for each layer would a labor-intensive and biased task. To avoid bias, a convolutional neural network founded on a U-Net architecture was employed for segmenting cross-sectional images of both the actin and nucleus channels. For training and validation, image sets were manually created by labeling each F-actin dot and nucleus border on the Y-Z cross-sectional images. The training set comprised 86 cross-section slices of the actin channel of Mesenchymal Stem Cells (MSCs), sourced from various image sets and microscopes and resized to 512×512 pixels. Out of the total images, 74 were used for training and 12 for validation. In the case of the cross-section nucleus, the training set included images from different slices and cells, with 9 randomly assigned to validation and 56 to training. The learning parameters were set at a learning rate of 0.001 and batch size of 1. The model underwent 200 epochs of training, accelerated by a Graphics Processing Unit (GPU). During training, the neural network works to minimize the loss function, emphasizing the pixel-to-pixel differences between the predicted and actual images. The loss function was adjusted to amplify the error for false-negative results for actin fibers to 20, a setting that was also used for the nucleus model.

To enable fiber assignment, fibers were labeled as apical or basal based on the mean Z coordinate of each fiber point. If the mean was higher than 40% of the nucleus height, the fiber was labeled as apical. If it was lower, it was tagged as a basal fiber. This 40% threshold is user-adjustable. We implemented a significant update to this part of the algorithm, deviating from the original published version.^28^ The modified algorithm considered the actual height of each nucleus rather than the layer count used before. This adjustment was essential for images that captured multiple nuclei of varying heights. It ensured the accurate assignment of apical or basal fibers based on the actual height of each nucleus, preventing any misclassification that may occur if layer count was used as the basis for fiber assignment.

#### Fiber and Nucleus Reconstruction (Fig. S1c)

As illustrated in Fig.1c, reconstructing individual F-actin structures began with the identified F-actin dots, referred to as contours from this point onward. Starting from the first layer, contours were connected across subsequent layers using the following rules: i) single overlap – if a contour overlapped only one contour from the previous layer, it was added to the same fiber; ii) one contour overlaps two from the previous layer – it was connected to the contour with the larger overlapping area; iii) no overlap – a new actin fiber was initiated.

After the reconstruction process, fibers with lengths less than 0.5 µm were discarded from the analysis. For each specific fiber within each cell in the dataset, measurements were taken for length, volume, and intensity. Branching information was integrated into the final statistics to indicate the number of points where a fiber split into multiple fibers. This additional data assisted in analyzing whether the actin structure was transitioning to a more net-like configuration. For nucleus reconstruction, the outlines of the nucleus on each Y-Z plane were combined to form a unified nucleus object. The volume of this object was calculated, and its dimensions were assessed by measuring the length of the rotated nucleus projection on the X-axis, the width on the Y-axis, and the height on the Z-axis.

### Statistical analysis of unbiased machine learning data

The probability distribution was used to anticipate how often specific variable values may occur. In this way, our initial analysis involved evaluating the performance of 41 distinct univariate probabilistic distributions to determine which one best captured the characteristics of the F-actin measurement data, thereby identifying the most appropriate distribution for modeling the observed F-actin (**Fig.S2&S3**) and nucleus (**Fig.S4**) features. For the probabilistic approach, “gamlss” package was used in R software 4.1.3 (R Core Team, 2022).^29^

During the trial of distributions, value type and ranges of measurements were considered. Therefore, i) 23 discrete (count) probability distributions were applied to basal fiber number, apical fiber number, basal branching node number, apical branching node number, and ii) 18 non-negative continuous probability distributions were applied to basal fiber length, apical fiber length, basal fiber volume, apical fiber volume, nucleus width, nucleus length, nucleus volume, and nucleus high. We used the method of maximum likelihood to estimate the parameters of all distributions so that the observed data were the most probable given the chosen distribution. The performance of the distributions were contrasted via Akaike’s information criteria (AIC)^30^ (**Table 1**).

**Table 1.**
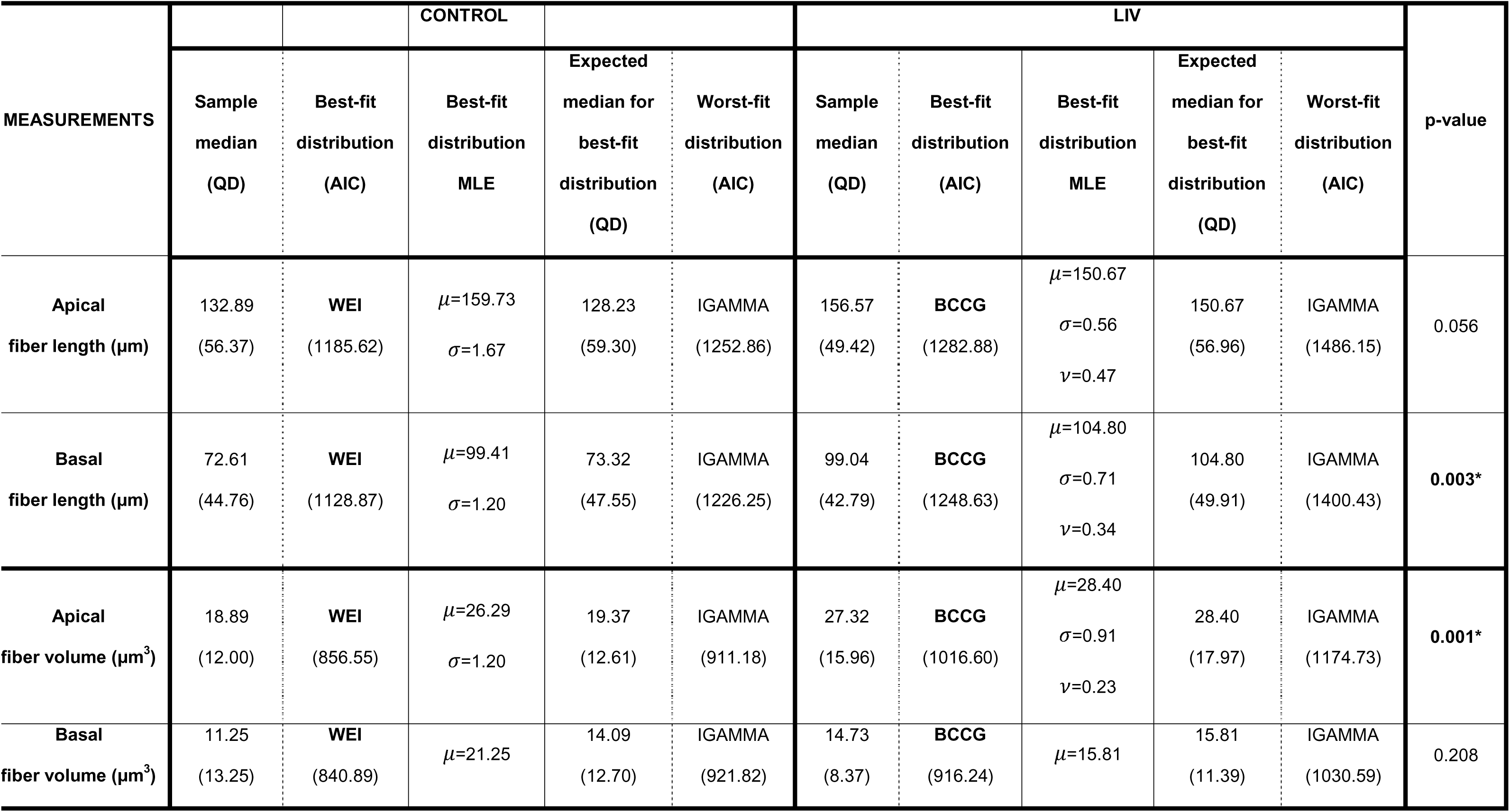
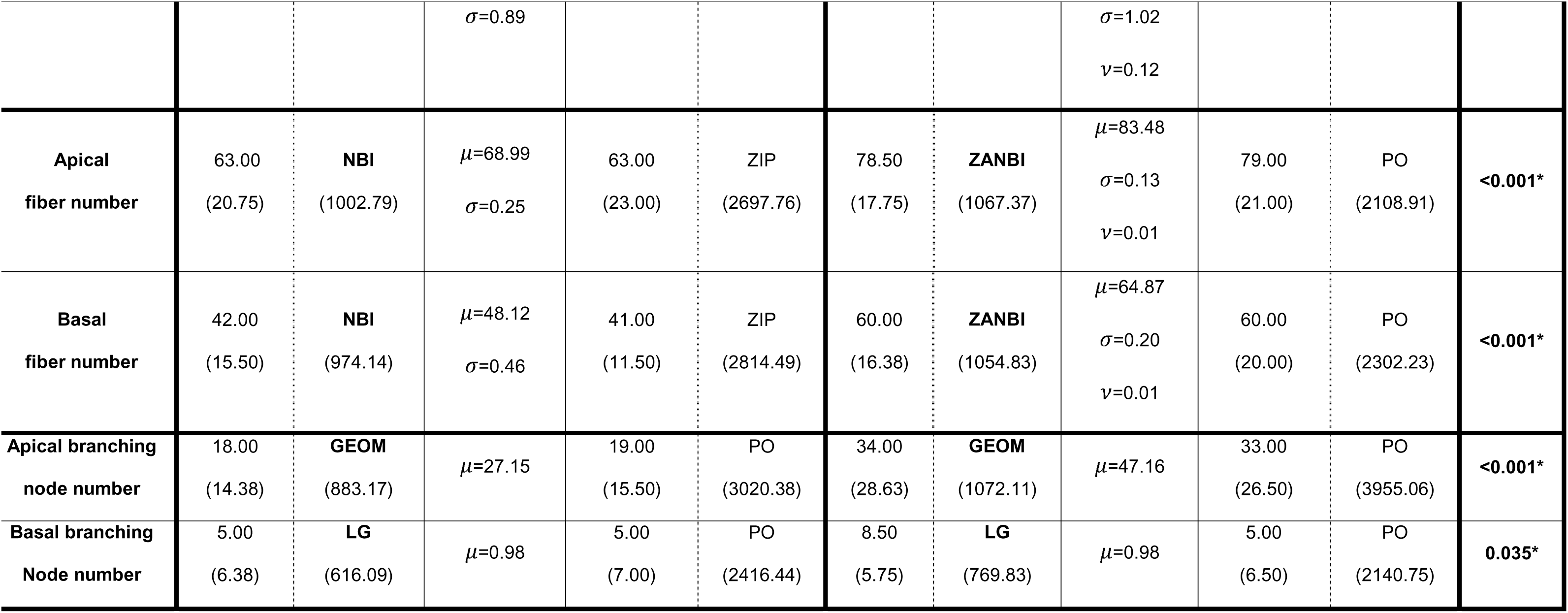
Maximum likelihood estimates and fitting performances of probability distributions. Bold indicates p-value<0.05. MLE - maximum likelihood estimation, QD - quartile deviation, AIC - Akaike information criteria.

As none of the measurements followed a normal distribution, median, and inter-quartiles were used for descriptive statistics (**Table 1**). Mann-Whitney U tests probed group differences and. Two-tailed p-values of less than 0.05 were considered statistically significant in all analyses. Probabilistic differences between the control and LIV groups, according to best-fit distributions, were presented in **Table 2**.

**Table 2.** Some specific probabilities of measurements according to best-fit distributions.

| MEASUREMENTS | CONTROL |  |  |  | LIV |  |  |  |
| --- | --- | --- | --- | --- | --- | --- | --- | --- |
|  | ≥50 | ≥100 | ≥150 | ≥200 | ≥50 | ≥100 | ≥150 | ≥200 |
| Apical<br>fiber length (μm) | 0.87 | 0.63 | 0.41 | 0.23 | 0.94 | 0.75 | 0.50 | 0.30 |
| Basal<br>fiber length (μm) | 0.65 | 0.37 | 0.19 | 0.09 | 0.84 | 0.57 | 0.29 | 0.11 |
| Apical<br>fiber volume (μm<sup>3</sup>) | 0.12 | 0.01 | <0.01 | <0.01 | 0.26 | 0.06 | 0.01 | <0.01 |
| Basal<br>fiber volume (μm<sup>3</sup>) | 0.11 | 0.02 | <0.01 | <0.01 | 0.11 | 0.02 | 0.01 | <0.01 |
| Apical<br>fiber number | 0.66 | 0.17 | 0.03 | <0.01 | 0.86 | 0.26 | 0.03 | <0.01 |
| Basal<br>fiber number | 0.38 | 0.08 | 0.01 | <0.01 | 0.64 | 0.12 | 0.01 | <0.01 |
| Apical branching<br>node number | 0.16 | 0.03 | <0.01 | <0.01 | 0.34 | 0.12 | 0.04 | 0.01 |
| Basal branching<br>Node number | 0.07 | 0.02 | <0.01 | <0.01 | 0.07 | 0.02 | 0.01 | <0.01 |

### Finite element model generation and analysis

To compare nuclear stresses between control and LIV conditions, three components were recreated *in silico*, nuclei, F-actin fibers, and cell plate. The initial unloaded nuclear geometry was modeled as an ellipsoid based on previously measured isolated nuclei images^25^ with volumes corresponding to experimental measurements (**Fig.2c**). The geometric surfaces were meshed using three-node shell, S3, elements, from which a volume mesh was constructed out of three-dimensional linear tetrahedron, C3D4, elements (**Fig. 5a**). C3D4 elements were used to mesh the nuclei because of their ability to match complex shapes. The initial nuclei model was composed of ∼18990 elements approximately 500nm in size. Each nucleus was assigned an linear elastic Elastic Modulus of 1.08 kPa and a Poisson’s ratio of 0.3.^25^

**Figure 2:**
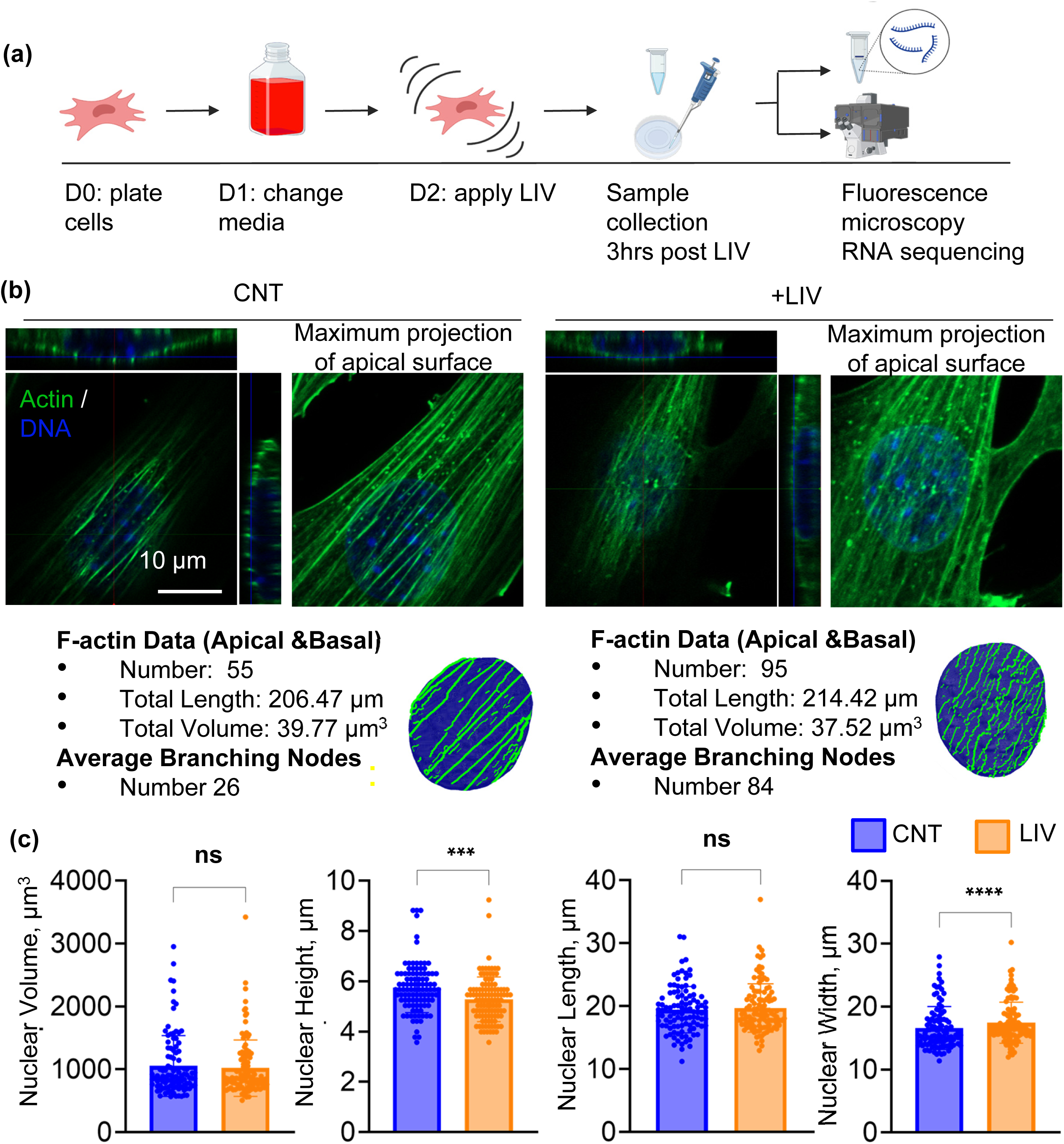
Low-Intensity Vibration (LIV) decreases nuclear height. **(a)** Experimental Workflow: Schematic outlining the experimental sequence. Mesenchymal Stem Cells (MSCs) are seeded on Day 0, transitioned to adipogenic medium on Day 1, and subjected to low-intensity vibrations (LIV) on Day 2. Cells are fixed with formaldehyde for imaging, and samples are collected for RNA sequencing three hours after the final vibration. **(b)** Top row: Confocal microscopy orthogonal views of actin fibers and nuclei. Left: a cell exposed to LIV; right: an untreated control cell. Bottom row: Corresponding 3D reconstructions and quantitative data of fiber and nucleus morphology, generated using the algorithm described in Figure S1. **(c)** Analysis of Nuclear Metrics: Bar graphs display mean values and standard deviations to compare nuclear characteristics between LIV-treated (+LIV) and control (-LIV) groups. Statistical analysis was performed in GraphPad Prism using the Mann-Whitney U test. No significant differences were found in volume and length (p=0.6613 and p=0.9232, respectively). Nucleus height shows a significant median decrease in the +LIV group (p=0.002), corresponding to a reduction of approximately 7.41% compared to controls. Nucleus width also shows a significant median increase in the +LIV group (p=0.0077), corresponding to an increase of approximately 6.85%, suggesting a potential compensatory widening due to the height reduction.

The cell modes for Control and LIV conditions included 9 equidistant and parallel aligned apical F-actin fibers spanning the x-axis (**Fig.5a**). The geometric surface of the actin stress fibers ran in tangency along the apical surface of the ellipsoidal nucleus. The fibers were 0.2 µm wide and distributed across 75% of the top half of the nuclei. The arced surface was meshed with, three-dimensional three-node membrane, M3D3, elements, assigned an analytical thickness value of 0.2 µm (**Fig.5a**). The end nodes of the membrane elements were then connected via three-dimensional two-node connector, CONN3D2, elements (**Fig.5a**). The cytoskeletal connector elements were used as axial force applicators, extending three times the nuclear length in x-axis, and converging with the base of the nuclei in the z-axis. (**Fig.5a**). Each beam was composed of ∼28 membrane elements, approximately 0.1μm in size, and four connector elements, two on each end. The actin stress fibers were modeled by two element types to achieve multifaceted properties. Membrane elements were utilized for their realistic contact behavior, while connector elements were used to realistically generate forces along the axis of the actin stress fibers. The parallel stress fibers were assigned mechanical properties representative of actin bundles with an elastic modulus 1000 Kpa and a Poisson’s ratio of 0.3.^31^

A cell culture plate at the base of initial nuclei was modeled as a simple rigid flat plate and meshed into a single linear hexahedral (brick), C3D8, element (**Fig.5a**). A C3D8 element was used to mesh the cell plate because the square geometry aligned with the desired part geometry. The square plate was 30μm x 30μm x 1μm in size, designed to be larger than the nuclei. The main role of the cell culture plate was to act as the models’ base boundary.

Unloaded nuclei models were first imported into Abaqus/ CAE 2021 for dynamic explicit simulations. The cell culture plate was modeled as rigid and constrained in all 6 degrees of freedom (DOF). Two contact conditions were used to define the model behavior, general contact and surface-to-surface contact. The general contact condition defined the connection between the nuclei and the cell culture plate. The contact interaction between these surfaces was designed to have relatively low friction (∼0.2), permitting minimal sliding between nuclear elements and the cell plate. The surface-to-surface contact condition was used to define the connection between the nuclei and the individual actin stress fibers. Surface-to-surface contact utilizes a primary-secondary definition, assigning either a balanced or pure weighting to the pair controlling which part governs deformation. The nucleus was defined as the primary surface in this relationship, as per recommended for contact conditions between continuum elements and membrane elements. The actin fiber membrane elements were meshed slightly finer than the nuclei in accordance with modeling recommendations for avoiding excessive penetration. The defined contact condition did however allow for slight penetration between the two surfaces. The contact interaction between the two surfaces was defined by a “rough” friction definition, hence after the model components came into contact, they became essentially conjoined. This functionality was desired to realistically simulate the behavior between apical actin and nuclei surface as reported^32^ as well as for avoiding challenges with membranes slipping off the nuclei during initial contact. Axial forces were applied to each F-actin fiber through the cytoskeletal connector beams attached to the stress fiber membrane elements. The forces were applied using a tabular amplitude, linearly increasing over time. An encastered boundary condition was applied to the end nodes of the cytoskeletal connector elements, constraining movement of the nodes along all 6 DOF. Each nuclei were pulled down until nuclear apex reached the experimentally measured nuclear height (**Fig.2c,** see supplementary movie 1), following the end of simulations, nuclear stresses were compared by extracting von Mises stress.^33^

## Results

### Small accelerations generate an out of phase nuclear motion in a LINC complex dependent manner

The lack of significant matrix deformations generated by LIV^34^ and the requirement for LINC-complex mediated nucleocytoskeletal connectivity to support LIV-induced mechanosignaling led us to consider an *inside-out* mechanism where LIV-induced dynamic accelerations cause the nucleus to oscillate out-of-phase relative to the cell body.^35^ To answer this question, we built an LIV bioreactor that can oscillate cells in a microscope chamber (**Fig.1a**). Acceleration output and voltage input relationship was linear, with a 0.7g, 90Hz LIV signal corresponding to approximately 6.5V (dashed red line, **Fig.1b**). We confirmed the acceleration reading by tracking the time-averaged travel path of fluorescent particles after setting the camera exposure to equal the wavelength of the input signal (**Fig.1c**). Lowering the exposure time to 1ms allowed us to track the motion of a single nucleus vibrating at 90Hz, 0.7g over 11 frames (**Fig.1d**). Reconstruction of the time displacement history showed a periodic motion. Close-grouping of time points at the positive and negative peaks indicate regions where the nucleus slows down before moving in the opposite direction (**Fig.1e**). Next, we labeled the cell body with Calcein AM to allow for tracking of both nuclei and cell edge together. Quantification of the phase shift across multiple cells (N=47) showed that, on average, the amplitude of nuclear motion matches that of the cell during a 90 Hz, 0.7 g vibration, inducing a total cell displacement of 22 µm. Simultaneous tracking of nuclei and cell edge revealed that nuclei move the same total distance as the cell edges but lag behind the edge during the phase when the highest accelerations occur: the nucleus lagged by approximately 0.2 ± 0.11 radians (∼0.35 ms delay), corresponding to an average peak displacement lag of 1.27 µm (17%).

Next, we tested the effects of disabling the LINC complex function during 90Hz, 0.7g vibrations. dnKASH cells were treated with +Dox to express mCherry-KASH. mCherry-KASH competes with the Nesprin-KASH domain binding, effectively disabling LINC connectivity (**Fig.1j**).^28,36^ Shown in **Fig.1k**, in LINC-disabled MSCs, the nuclear phase lag was doubled (p<0.05), suggesting that relative nuclear motion is in-part regulated by LINC connectivity. We then tested the dependence of phase lag on LIV frequency, increasing the frequency from 90Hz to 180Hz and retaining a 0.7g acceleration. No correlation between frequency and phase lag was found (**Fig.S2a**) while increasing the acceleration from 0.5 to 1.1g at 90Hz increased the phase lag (**Fig. S2b,** p<0.05).

### Small accelerations cause accrual of apical F-actin fibers with larger cross-sections, and increase in actin and focal adhesion-related genes

Our recent findings indicate that LIV does not alter microtubule dynamics^37^, leading to our focus on regulation of F-actin. To quantify F-actin changes, we utilized a novel machine learning based methodology for accurate reconstruction of nuclei-associated F-actin, quantifying individual F-actin features such as length, cross-section and volume from microscope images (**Fig.S1**).^28^ MSCs were fixed for 3h after a 7h application of 0.7g, 90Hz LIV (six repeats of 20min ON, 60 min OFF) and imaged via a Zeiss LSM 900 confocal microscope (**Fig.2a,** n=110 cells/grp). Shown in **Fig.2b**, cells subject to LIV had enhanced connectivity of the F-actin network with an increased number of smaller fibers connecting larger ones. A representative reconstruction from control (CNT) and LIV groups demonstrates that small accelerations a robust increase—by approximately one-third—in fiber number and a threefold increase in branching points. Compared to CNT, cells subject to LIV did not show changes in nuclear volume or length, but did show a 7% (p<0.001) reduction in nuclear height and a 7% increase in width (p<0.0001), consistent with nuclear flattening (**Fig.2c**).

We analyzed the aggregated F-actin features by pooling all actin data together. Depicted in **Fig. 3b**, cells exposed to LIV displayed a 24% increase in apical (p<0.001) and a 43% (p<0.001) increase in basal F-actin fiber numbers. We also measured branching points, defined as points where individual F-actin fibers intersect.^28^ Basal fibers, on average, had 3-fold fewer branching points compared to apical fibers; LIV induced only a modest 7% increase in basal fiber branching (p<0.05). In contrast, LIV increased the apical fiber branching by 89% (p<0.001, **Fig.2b**). No changes in apical or basal fiber lengths were found (**Fig.3c**). Consistent with increased apical F-actin connectivity, cross-section and volume of apical F-actin fibers were 20% larger after LIV exposure (p<0.0001) when compared to controls. Basal fiber thickness and volume decreased by 5% (p<0.0001) and 11%(p<0.001), respectively (**Fig.3d & 3e**, p<0.001).

**Figure 3:**
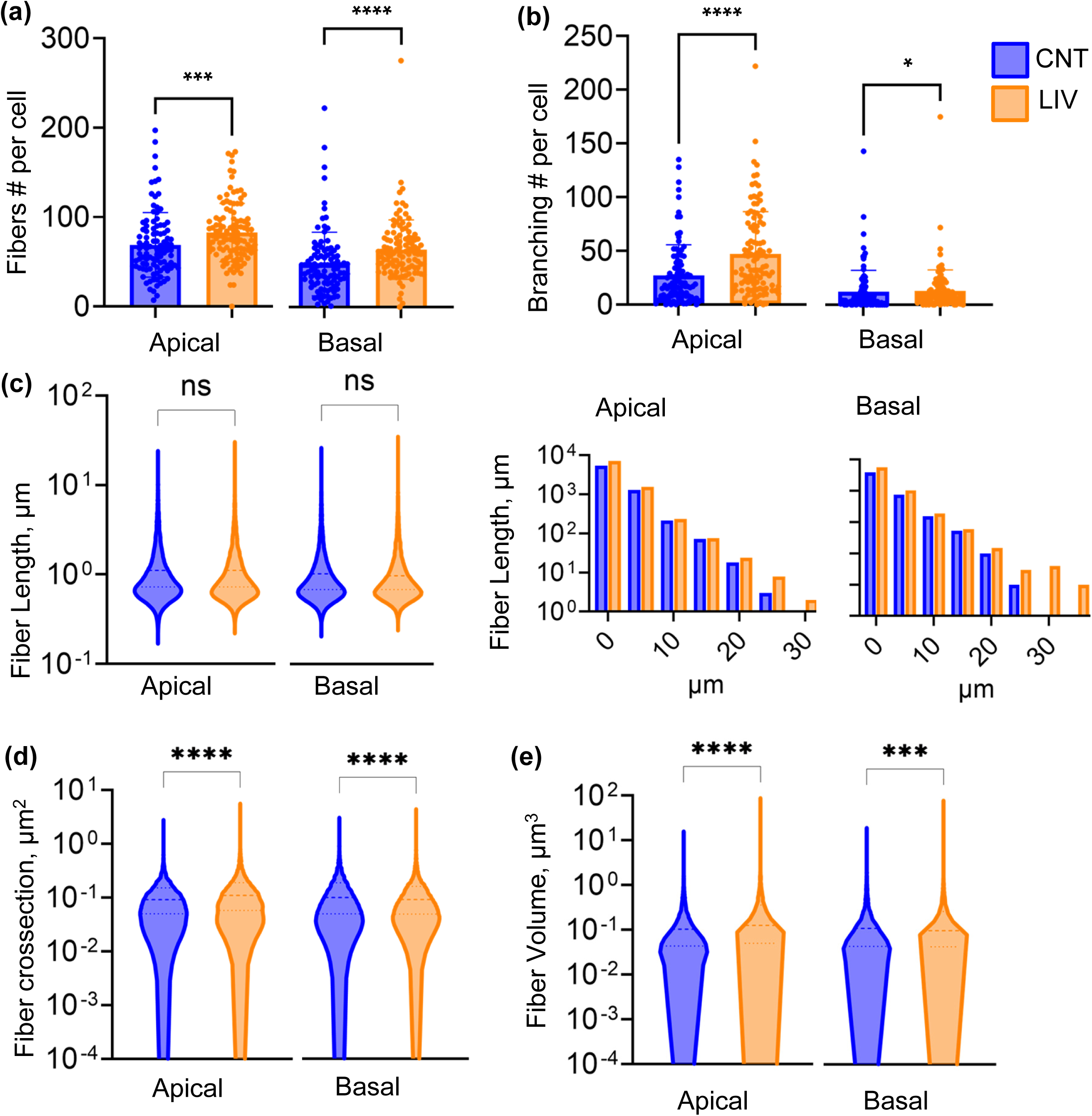
Quantitative Analysis of Actin Fiber Metrics under LIV and Control Conditions. **(a)** Number of Actin Stress Fibers per Cell: Compares the median number of apical and basal actin fibers per cell in LIV-treated (+LIV, n=110 cells) and control groups (-LIV, n=102 cells) using Mann-Whitney tests. Significant increases were observed in the median number of apical fibers (+LIV=78.5, -LIV=63.0 p=0.0004; 24.6% increase) and basal fibers (+LIV=60, -LIV=42; p<0.0001; 42.86% increase). **(b)** Number of Branching Points per Cell: Compares the median number of branching points, areas where a fiber divides to form a net-like pattern, between +LIV (n=110 cells) and -LIV (n=102 cells) groups. Significant differences were observed for both apical fibers (median +LIV=34, -LIV=18; p<0.0001; 88.89% increase) and basal fibers (median +LIV=8.5, -LIV=5; p=0.0350; 7% increase). Increased branching in the +LIV group suggests enhanced fiber network complexity. **(c)** Fiber Length Distribution: The left graph is a violin plot aggregating individual fiber lengths from 110 cells in the +LIV group and 102 cells in the -LIV group. No significant differences in median fiber length for both apical (+LIV n=9100 fibers, -LIV n=7037 fibers; p=0.0567) and basal fibers (+LIV n=7071 fibers, -LIV n=4908 fibers; p=0.1111) were observed. The right graph is a histogram showing the fiber length distribution, suggesting an increased number of longer fibers (>25 μm) in the +LIV group. **(d)** Individual Fiber Thickness: Reports changes in median fiber thickness based on aggregated individual fiber measurements from +LIV (apical n=9100 fibers, basal n=7071 fibers) and -LIV (apical n=7037 fibers, basal n=4908 fibers) groups. While apical fibers showed a significant increase in thickness (+LIV median=0.1096 μm², -LIV median=0.09142 μm²; p<0.0001; 19.88% increase), basal fibers demonstrated a significant reduction in thickness (+LIV median=0.9698 μm², -LIV median=1.018 μm²; p<0.0001; 4.72% decrease). **(e)** Individual Fiber Volume: Presents volume data based on individual fibers aggregated from +LIV (apical n=9100 fibers, basal n=7071 fibers) and -LIV (apical n=7037 fibers, basal n=4908 fibers) groups. Significant increases were noted in apical fiber volume (+LIV median=0.1247 μm³, -LIV median=0.1032 μm³; p<0.0001; 20.83% increase), while a significant decrease was observed in basal fiber volume (+LIV median=0.09678 μm³, - LIV median=0.1089 μm³; p=0.0008; 11.14% decrease). All data shown in panels (a–e) were derived from 3D reconstructions using the algorithm detailed in Figure S1. Statistical tests were performed in GraphPad Prism.

We then asked if there were gene expression changes concomitant with LIV-induced structural changes. Differentially expressed (p<0.05) genes 3h after LIV exposure (n=3/grp) were identified via DESEQ2 analysis. Hierarchical heat maps of differentially expressed genes showed a clustering of LIV treatments (**Fig.4a**) with a total of 372 genes (218 up, 154 down) displaying significant (p<0.05) differences in transcriptional activity between CNT and LIV groups (**Fig.4b**). DAVID Pathway analysis showed significant enrichment (p<0.05) in 36 categories for upregulated genes (**Table.S2)** and 42 categories for downregulated genes (**Table.S3**). Upregulated categories included actin filament, cilium assembly, fluid shear stress and atherosclerosis with focal adhesions being the most upregulated category, containing focal adhesion and actin regulators such as Fyn^38^ and Pak3, integrin beta 3 and Zyxin^39^(**Fig.4d**).

**Figure 4:**
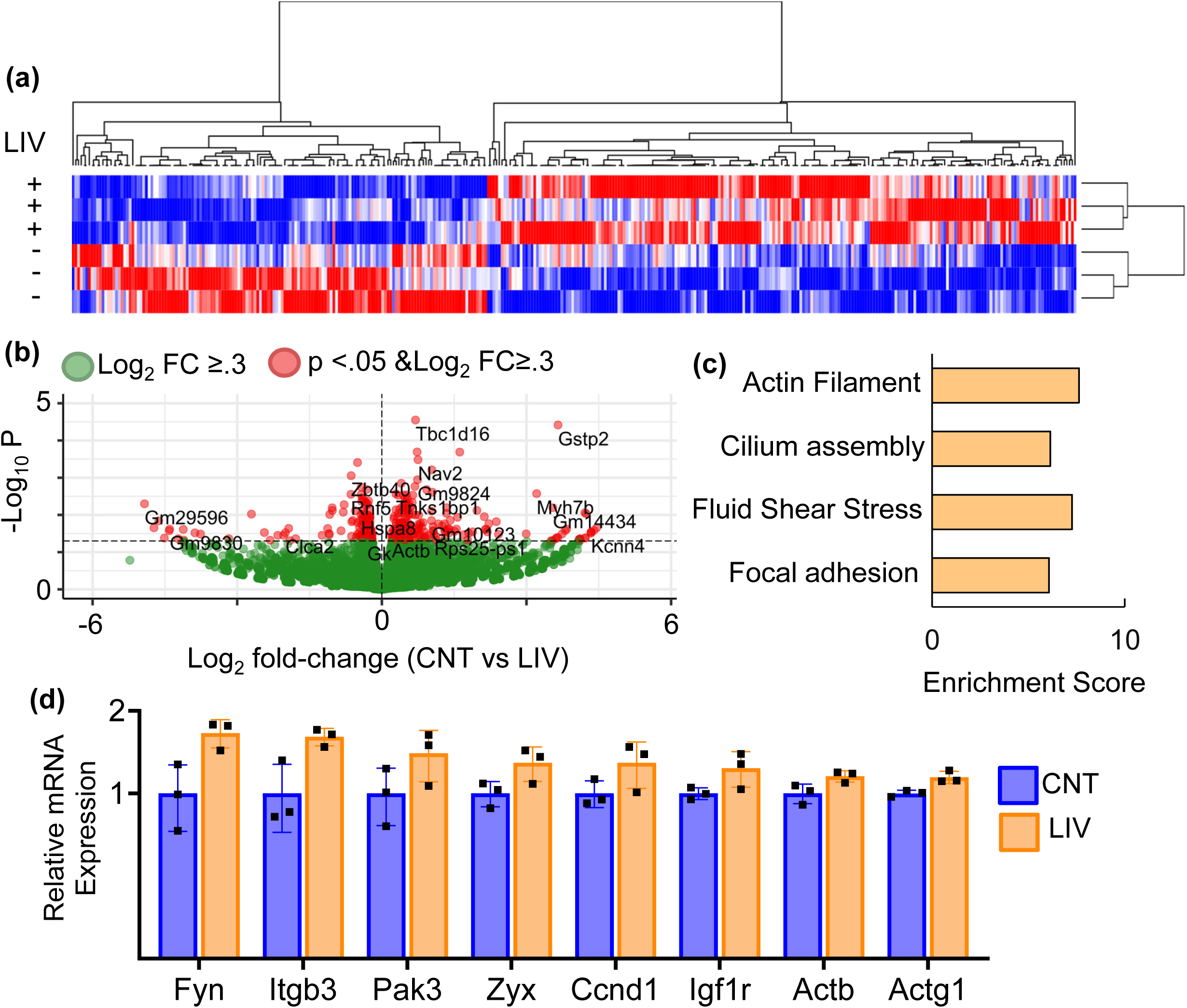
LIV increases mRNA expression of actin and focal adhesion-related genes. (a) Differentially expressed (p< 0.05) genes 3hr after LIV exposure (n=3/grp) were identified via DESEQ2 analysis. Hierarchical heat maps of differentially expressed genes showed a clustering of +LIV treatments with a total of 372 genes (218 up, 154 down) between CNT and LIV groups with p<0.05 statistical significance. **(b)** (DAVID Pathway analysis showed significant enrichment (p<0.05) on 78 (**Table.S3 and S4)**. **(c)** Upregulated categories included Actin filament, cilium assembly, Fluid shear stress, and athelerochlorosis with Focal adhesions being the most upregulated category. **(d)** This category included focal adhesion and actin regulators such as Fynand Pak3, integrin beta 3 and Zyxin.

### Small accelerations result in a distinct nuclei-associated F-actin architecture and increases in nuclear stress

As aggregated statistics revealed significant nuclei-associated F-actin alterations, we next asked whether CNT and LIV treated cells have different distributions of F-actin and nuclei features. Shown in **Fig. S3**, out of 41 possible statistical distributions tested, apical and basal fiber lengths and volumes of all +LIV treated groups were best represented by Box-Cox Cole and Green distribution (BCCG) while all CNT groups were represented by Weibull (WEI) distribution. Cell based distributions (**Table 1**) showed a 44% increase in the apical fiber volume (p<0.001) with no change in apical fiber length indicating thicker fibers while basal fibers were 36% longer (p<0.01) with no changes in fiber volume. Discretely distributed fiber and branching fiber data (**Fig. S4**) showed that apical and basal fiber numbers of +LIV treated cells were best represented by a Negative Binomial type I (NBI) distribution while the CNT group was best fitted to a zero adjusted negative binomial (ZANBI) distribution. LIV increased both apical and basal fiber per cell by 30% (p<0.001) and 42% (p<0.001), respectively. Apical fibers showed 88% more branching points (18 in CNT as compared to 34 in LIV, p<0.05), while the average number of basal fibers per cell was 5 in CNT cells, with LIV increasing the number to 8.5 per cell (p<0.05). Nuclei features between CNT and LIV showed the same statistical distributions (**Fig. S5**).

We next sought to develop generalized models to identify the F-actin features that differentiate LIV from control cells. We assumed that only longer (>4µm) F-actin fibers spanning the apical nuclear surface contribute to nuclear shape. Utilizing bootstrap confidence intervals based on NBI distributions of apical fibers, we estimated that there were 9.4 ± 6.5 and 9.6 ± 6.9 apical fibers in CNT and LIV conditions, respectively (**Fig.S6**). As detailed in the methods section, FE models sought to capture the apical F-actin and nuclei features (**Fig. 5a**) and compared the forces required to deform the nuclei to generalized shapes They also quantified the mechanical stresses generated inside the nuclei by applying F-actin forces to an undeformed nuclei (**Supplementary movie 1**). Shown in **Fig.5b**, the FE model identified the nuclear surface directly under load bearing actin fibers and mid-nuclei as locations of peak stress. Assuming no changes to the nuclear stiffness, FE predicted that 30% greater total F-actin force was required to deform LIV-treated nuclei on the apical nuclear surface (117nN) when compared to CNT nuclei (90nN). As a result, LIV-treated nuclei had a greater number of elements with higher Von Mises Stress, increasing the average stress by 18% (p< 0.0001), from 0.00043 MPa (0.43kPa) to 0.00052MPa (0.52kPa). As we previously reported that LIV increases nuclear stiffness,^25^ implementing stiffer nuclei in our LIV model (increased to 1.85 kPa from 1.08 kPa, **Fig.S7**) predicted a 50% increase in F-actin force, a 84% increase in average intra-nuclear Von Mises Stress (p<0.0001), thereby identifying nuclear mechanical properties as a key driver of nuclear stress.

**Figure 5:**
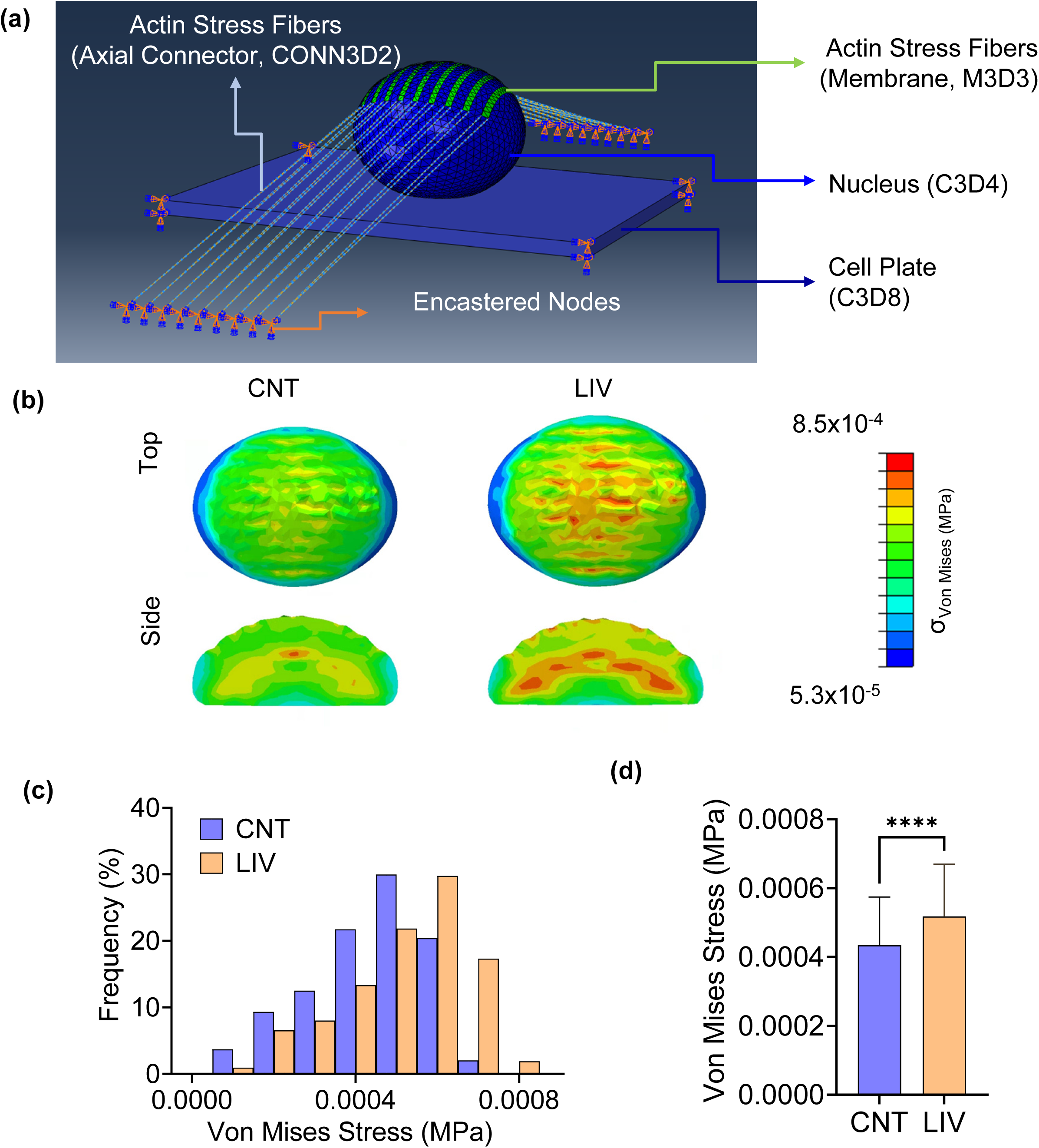
LIV increases nuclear stress. **(a)** We generated FE models to capture the apical F-actin and nuclear features that best represent both CNT and LIV conditions based on statistical data. We assumed that only longer (>4µm) F-actin fibers that span apical nuclear surface would contribute to the nuclear shape. Utilizing bootstrap confidence interval based on NBI distributions of apical fibers, we estimated 9.4 ± 6.5 and 9.6 ± 6.9 apical fibers per cell for CNT and LIV conditions, respectively. Our FE model identified the nuclear surface directly under load bearing actin fibers and mid-nuclei as locations of peak stress. **(b)** Assuming no changes to nuclear stiffness, the FE model predicted that 30% more total F-actin force is required to deform LIV treated nuclei on the apical nuclear surface (117nN) when compared to CNT nuclei (90nN). **(c)** LIV treated nuclei had more elements with higher Von Mises Stress **(d)** increasing the averaged stress by 18% (p< 0.0001) from 0.00043 MPa (0.43kPa) to 0.00052MPa (0.52kPa).

## Conclusions and Discussion

Mechanical signals are critical for cell-mediated regulation of tissue form and function. Traditionally, studies have focused on substrate strain and cell distortion as primary mechanisms for transmitting mechanical information to the cell. In contrast, our findings suggest that acceleration in and by itself can serve to drive cellular adaptation. This ability to recognize a very small physical signal driven by the relative motion of the nucleus with respect to the cell membrane, and autoregulated by the degree of nuclear actin tethering, is predicated to influence cell outcomes in adherent, suspended, and embedded cells

In testing the hypothesis that the nucleus moves relative to the cell body during even small accelerations, we found that the nucleus exhibits a phase shift relative to the cell body, a shift that is dependent both on the acceleration intensity as well as the degree to which the nucleus is tethered to the membrane. This phase shift induces changes in actin architecture, and further autoregulation of the mechanical signal that influences transcriptional machinery. The magnitude of the measured relative motion due to LIV was, on average, about 17% of the total cell motion amplitude (1.27µm, **Fig.1h**). This is comparable to previous FE models^35^ that predicted relative nuclear motion of nucleus between 0.7 µm and 1.8 µm for a 100Hz, 1g LIV signal as well as the recent experimental findings in HeLa cells.^40^ While increasing frequency from 90Hz to 120Hz slightly increased nuclear phase shift, further increases in frequency resulted in a decreased phase shift. As such, we found no correlation between frequency and nuclear phase shift (**Fig. S2a**). On the other hand, increases of acceleration magnitude at 90Hz showed a positive correlation with the nuclear phase shift (**Fig.S2b**, p<0.05). The correlation of phase shift with acceleration, but not with frequency, suggests that cells may exhibit distinct resonance frequencies in a culture dish based on various factors, including cell morphology, nuclear size and cytoskeletal configuration. Extrapolating this to an *in vivo* context, the dynamics of functional activities such as swimming, running or flying would generate widely varying cell distortions depending on the forces causing strain of the substrate; i.e., an osteoblast on the periosteal surface of bone versus an MSC suspended in marrow. Still, every cell in the body experiences the same overall acceleration and deceleration to which the organism is subject to, and would ‘tune’ the perceived signal based on membrane-to-nucleus tethering, creating a ubiquitous, and simple means for any cell to regulate the perceived mechanical environment of the nucleus.

Importantly, we showed that disabling LINC complex doubles the nuclear motion (**Fig.1k**). This suggests that nucleo-cytoskeletal configuration plays a major role in regulating nuclear motion. While this may indicate that increased nuclear motion results in larger forces, we previously reported that LIV-induced increases in nuclear stiffness do not occur when LINC-complex function is disabled.^41^ Thus, nuclei may not directly sense acceleration per se, but rather, nuclear motion serves as a sensing mechanism as the nucleus tugs at the actin cytoskeleton, generating *inside-out* forces.^41^ LINC-complex dependent activation of RhoA signaling at focal adhesions^22^ and the AFM-measured apparent stiffness of F-actin fibers post-LIV^23^ further supports this “inside-out” mechanism where forces are generated via nuclear tugging on the cytoskeletal elements. In this fashion, the level of nucleo-cytoskeletal coupling may regulate forces generated inside cells in response to dynamic accelerations, and further that variations in nucleo-cytoskeletal tethering (e.g., simulated microgravity^42^ or cell type) would modulate the degree of mechanical input the cell experiences. Here, we performed the first structural quantification of nuclei-associated F-actin architecture in response LIV. Unbiased and data-driven reconstruction of nuclei-associated F-actin networks showed that LIV-induced F-actin networks were statistically distinct from control conditions and that LIV increased number of branching points and number of small fibers (<4µm) on the apical nuclear surface, indicating an increasingly connected apical network in response to LIV-induced nuclear motions.

While the nucleus however, is mechanosensitive to direct displacement as low as 0.3µm on the nuclear surface,^43^ applying substrate stretch at sufficiently high levels (10%) leads to nuclear softening to protect the genome.^44,45^ Similarly, 4% stretch is also sufficient to decouple nuclei from the cytoskeleton to protect the cell nucleus from damage.^46^ Instead of decoupling the nuclei, LIV both increases the nuclear stiffness in a LINC complex dependent manner^25^ and increased F-actin accumulation around nuclei. This suggests that LIV exerts smaller forces on cell nuclei when compared to substrate stretch.

Our findings further show that F-actin fibers had a larger volume and thicker cross-section resulting in a flatter nucleus, indicating higher intra-nuclear stresses in LIV treated cells. These findings, in conjunction with a total of 372 differentially expressed genes between LIV and control conditions, indicate that F-actin-mediated forces both within and around the nucleus leads to intranuclear organization changes to affect gene expression. For example, our recent findings demonstrate that disabling LINC mediated forces in the cell nuclei was sufficient to reduce osteogenic^47^ as well as adipogenic^48^ differentiation via effecting heterochromatin dynamics. FE model predicted that 30% more total F-actin force is required to deform the LIV treated nuclei on the apical nuclear surface (117nN) when compared to control nuclei (90nN). While our predicted force values were not experimentally validated, the average force per F-actin fiber to maintain nuclear shape was between 10 and 13nN. This is comparable to experimental estimation of 10 to 67nN forces applied to apical nuclear surface by a single F-actin fiber in smooth muscle cells.^49^ Similarly, total force applied by actin fibers on the actin cap is comparable to 100nN cell-cell contacts measured in epithelial cells.^50^ FE model findings further indicated an 18% increase of intra-nuclear stress increases showing that forces experienced by cells under LIV are comparable to forces within cell mechanical environment.

A limitation of our model is the homogenous stiffness of nuclei, which predicted increased stress at the nuclear interior. Having stiffer heterochromatic and softer euchromatic regions, nuclei experience non-homogenous deformations.^51^ We previously established a model that can capture the mechanical heterogeneity of the cell nuclei which showed that the nuclear envelope acts to shield nuclei interior from high stresses.^52^ Therefore improving intra-nuclear stress predictions of this model is required to better model changes in the nuclear structure.

In summary, we have shown that accelerations, less than 1g and similar to those engendered by terrestrial locomotion, induce nuclear motion in a LINC-complex dependent manner, in turn augmenting the F-actin cytoskeleton and increases mechanical stress on cell nuclei. Further identification of nuclear motion, as a means to sense and respond to exogenous mechanical signals, achieved independent of substrate strain, should help establishing a causal understanding of how acceleration affects cell identity. Such information has the potential to provide mechanistic guidelines on how to use mechanical signals such as LIV for clinical and regenerative approaches to improve cell anabolism.

## Acknowledgments

This study was supported by AG059923, AR075803, AR075314, P20GM109095, NSF1929188, NSF 2025505, and NSF 2431083

## Data Availability

Software code and packaged release information. All RNA-seq data is provided as supplementary data.

## Competing interests

Clinton R. Rubin has several issued and pending patents on the use of low intensity vibration and extremely low magnitude mechanical signals to promote cell proliferation and differentiation, both in vitro and in vivo. He is also the founder of Lahara Bio and Marodyne Medical. No other author(s) declare no competing interests, financial or otherwise.

## Contributions

Nina Nikitina: concept/design, data analysis/interpretation, manuscript writing

Jonathan Wadsworth: concept/design, data analysis/interpretation, manuscript writing

Nurbanu Bursa: data analysis/interpretation, manuscript writing

Madison Goldfeldt: concept/design, data analysis/interpretation, manuscript writing

Chase Crandall: data analysis, final approval of manuscript

Sean Howard: data analysis, final approval of manuscript

Amevi Semodji: data analysis, final approval of manuscript

Anamaria Zavala: data analysis, final approval of manuscript

Stefan Judex: data interpretation, manuscript writing, final approval of manuscript

Janet Rubin: Judex: data interpretation, manuscript writing, final approval of manuscript

Trevor J Lujan: data analysis, final approval of manuscript

Clare K Fitzpatrick: data analysis, final approval of manuscript

Clinton R. Rubin: data interpretation, manuscript writing, final approval of manuscript

Aykut Satici: concept/design, data analysis/interpretation, financial support, final approval of manuscript

Gunes Uzer: concept/design, data analysis/interpretation, financial support, manuscript writing, final approval of manuscript

## SUPPLEMENTARY MATERIAL

**Funding support:** AG059923, AR075803, P20GM109095, NSF1929188, NSF 2025505 and NSF 2431083

**Figure S1:**
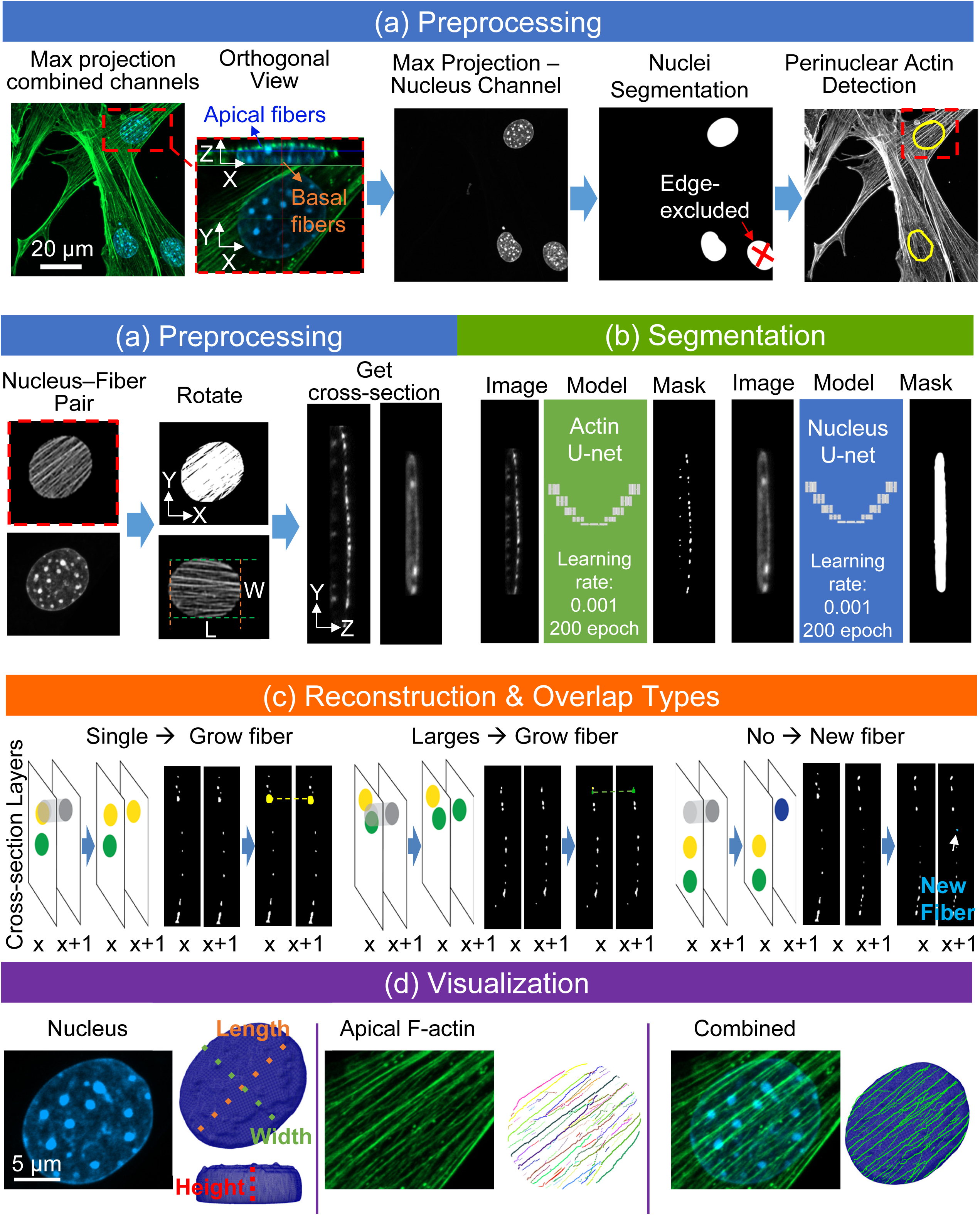
Stress Fiber Reconstruction Algorithm: **(a)** *<u>Preprocessing</u>:* maximal projections of the nuclear channel were segmented to detect nuclei and define ROIs. Each nucleus–fiber instance was cropped and rotated in the X–Y plane to align dominant fiber orientation using Hough Transform. Rotated stacks were converted into Y-Z cross-sections. **(b)** *<u>Segmentation</u>*process for actin fibers and nuclei in Y-Z cross-sectional confocal images using two distinctly trained U-Net CNN models. Training details for the neural network. The learning parameters used were a learning rate of 0.001, a batch size of 1, and 200 epochs. The loss function, which quantifies the differences between predicted and target images, was minimized during training. To prioritize false-negative results the loss function was adjusted to increase the error by 20 for both models **(c)** Actin fiber *<u>reconstruction</u>*process based on contour intersections across successive cross-sectional layers of actin masks. In this process, contours of detected F-actin blobs on cross-sectional masks are extended from the initial layer. These contours are methodically connected to corresponding ones on subsequent layers, following specific overlap criteria: 1. Straightforward continuation: A single contour from layer x+1 overlaps exclusively with one contour from layer x. This indicates a direct continuation of the actin fiber. 2. Overlap with multiple contours: When a contour from layer x+1 overlaps with several contours from the previous layer x, it is integrated into the fiber with the contour that has the largest overlap area. This scenario often represents a branching point. 3. Formation of a new actin fiber: If a contour on layer x does not overlap with any contours from the preceding layer x+1, a new actin fiber object is formed, signifying the start of a separate fiber. **(d)** A side-by-side comparison of original confocal microscopy images and their respective 3D reconstructions. The first column shows the nucleus and its mesh reconstruction. The middle column focuses on actin fibers, differentiated by color in the reconstruction to depict individual fiber paths. The last column displays the actin fibers superimposed on the nucleus.

**Figure S2:**
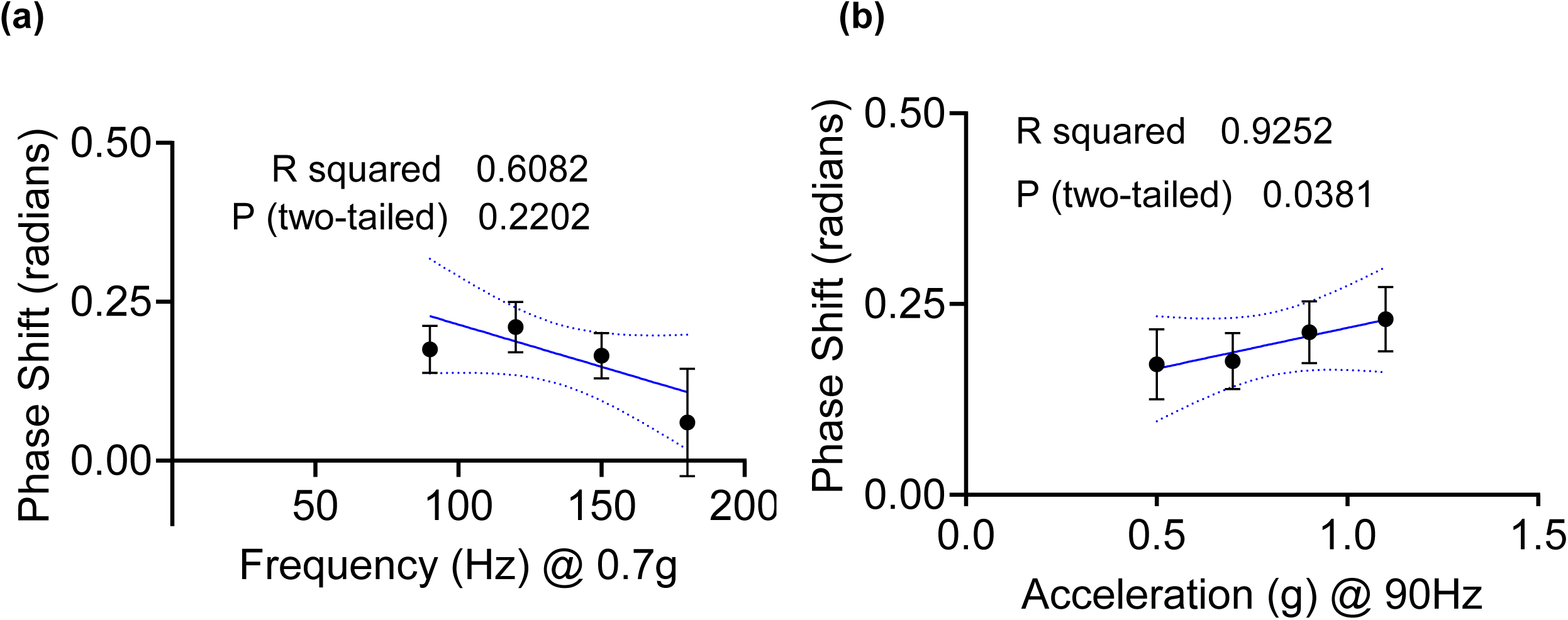
Nuclear phase shift correlates with acceleration magnitude and total cell displacement but not with frequency. Simple linear correlation analysis showed that **(b)** when acceleration was kept contact at 0.7g, increasing frequency did not result in a significant correlation with measured phase shift values. **(b)** When frequency was kept at 90Hz, nuclear phase shift increased with increased acceleration (P<0.05).

**Figure S3:**
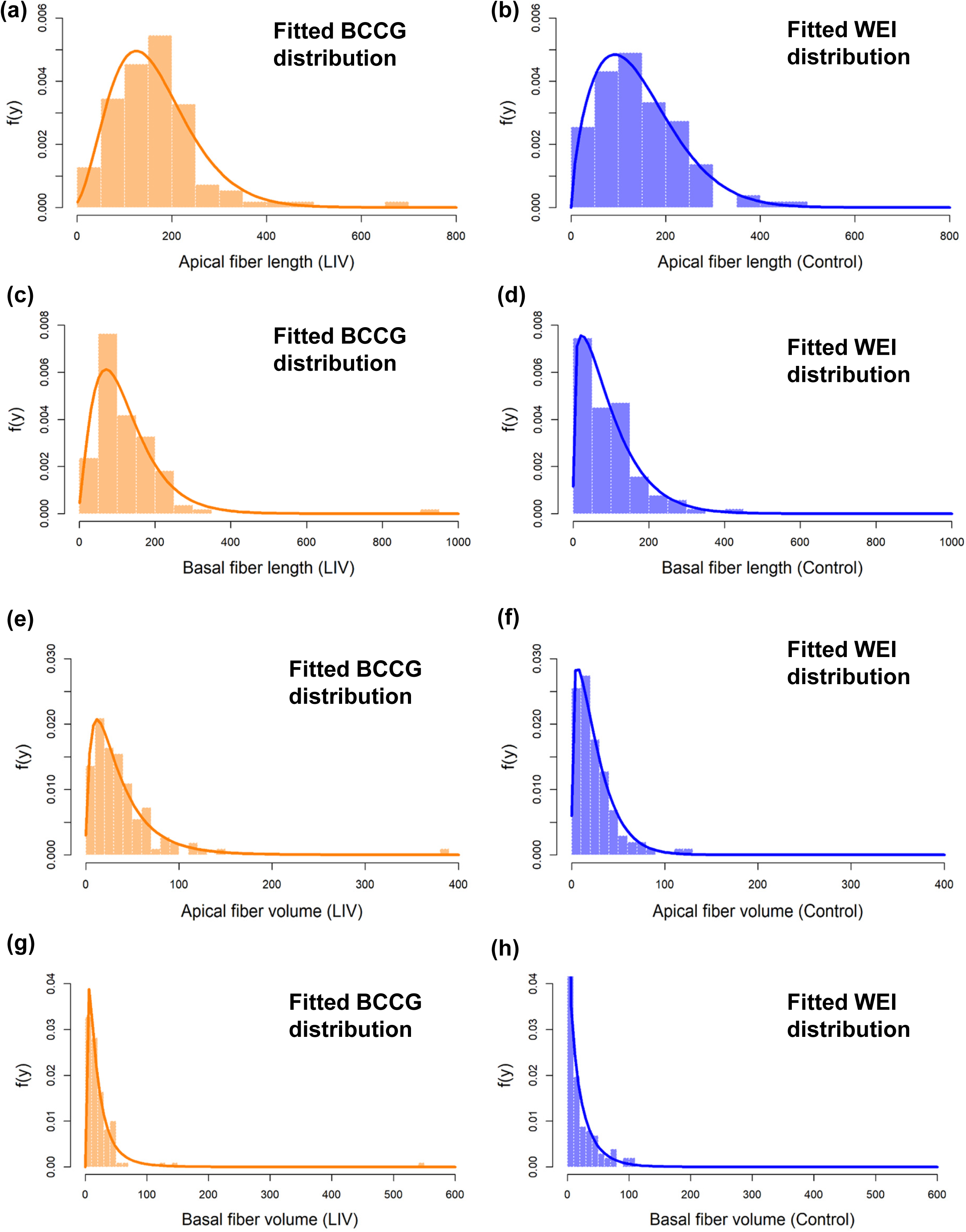
Fitting performance of best distributions for fiber length and volume.

**Figure S4:**
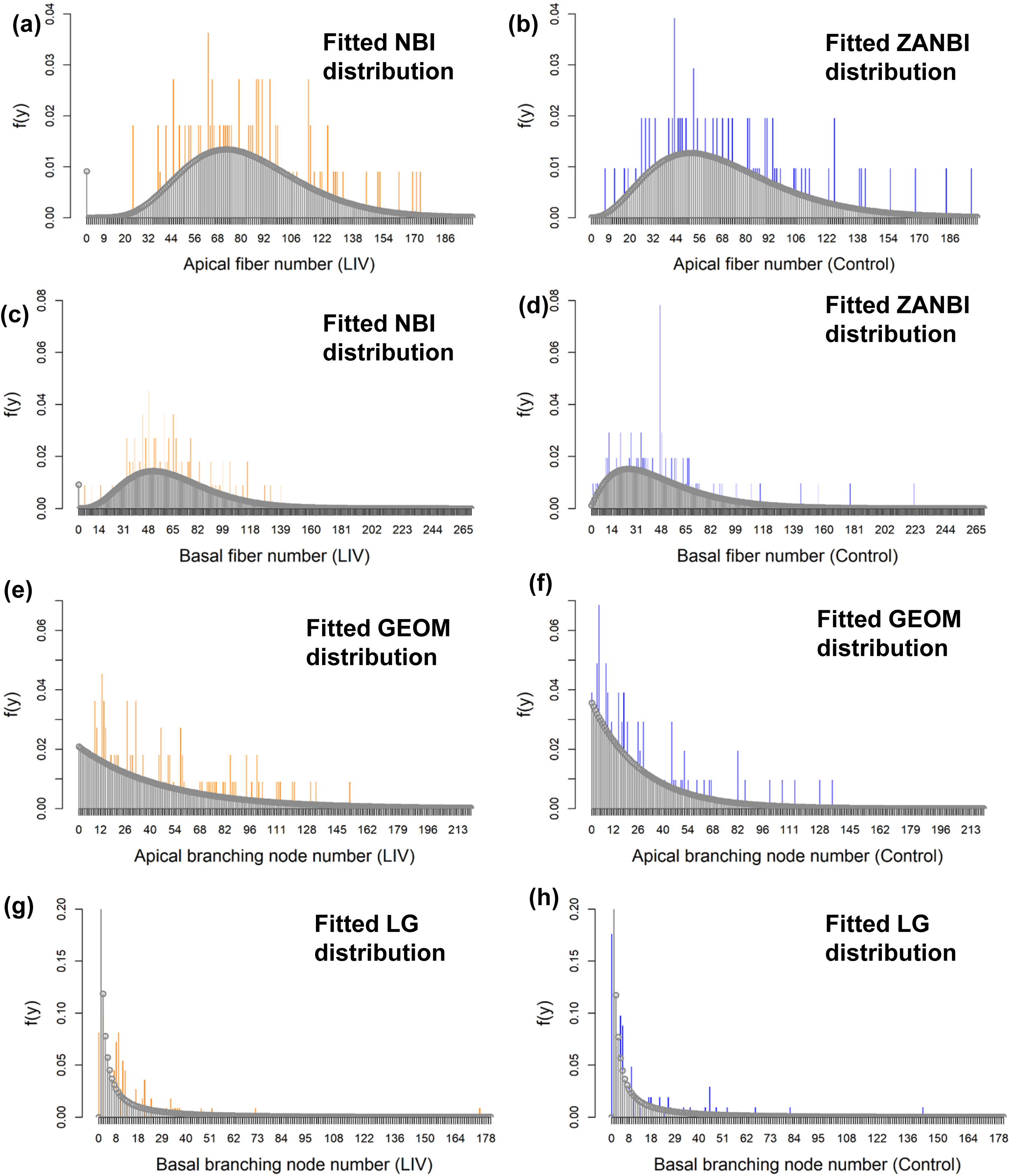
Fitting performance of best distributions for fiber number and branching number.

**Figure S5:**
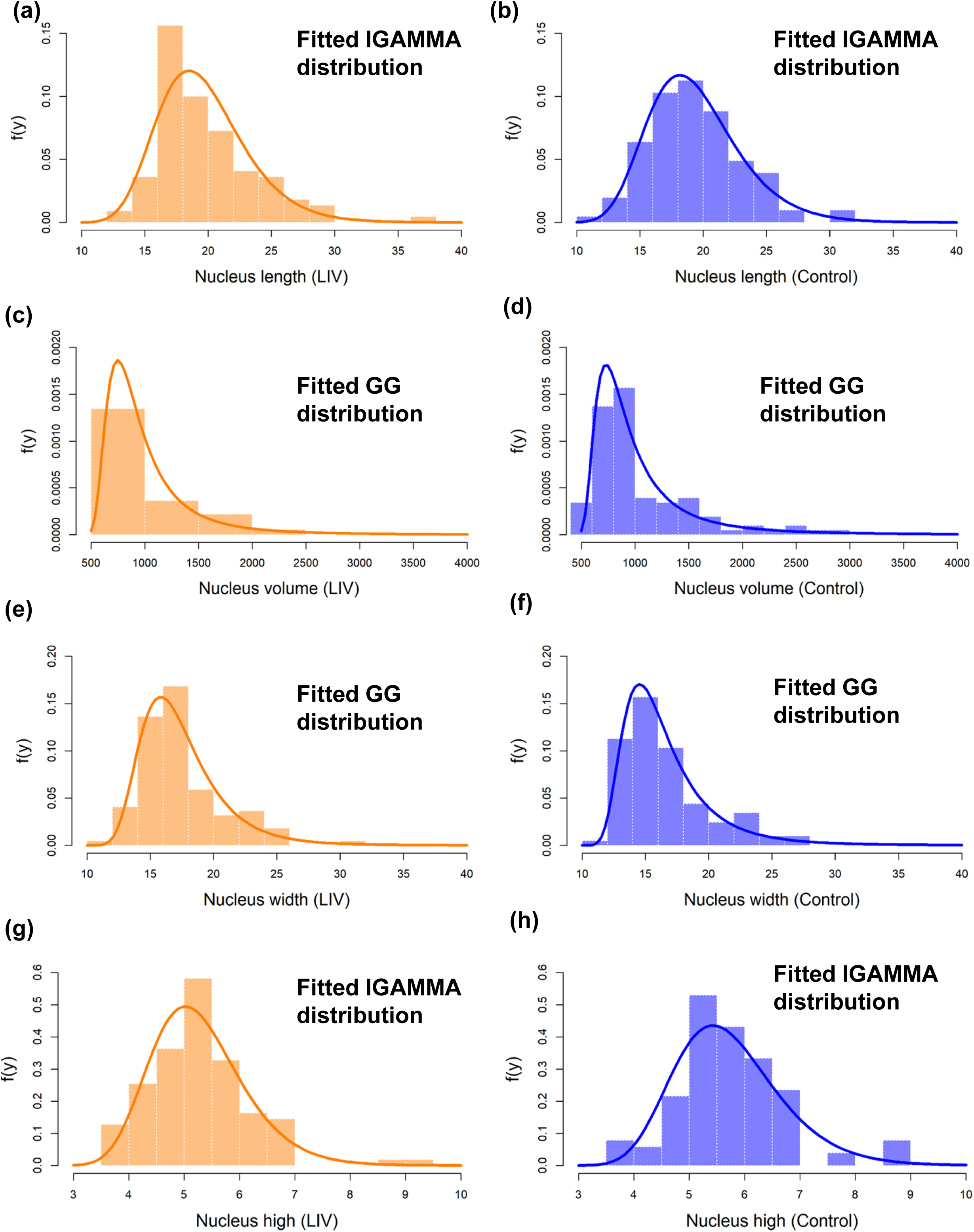
Fitting performance of best distributions for nucleus measurements.

**Figure S6.**
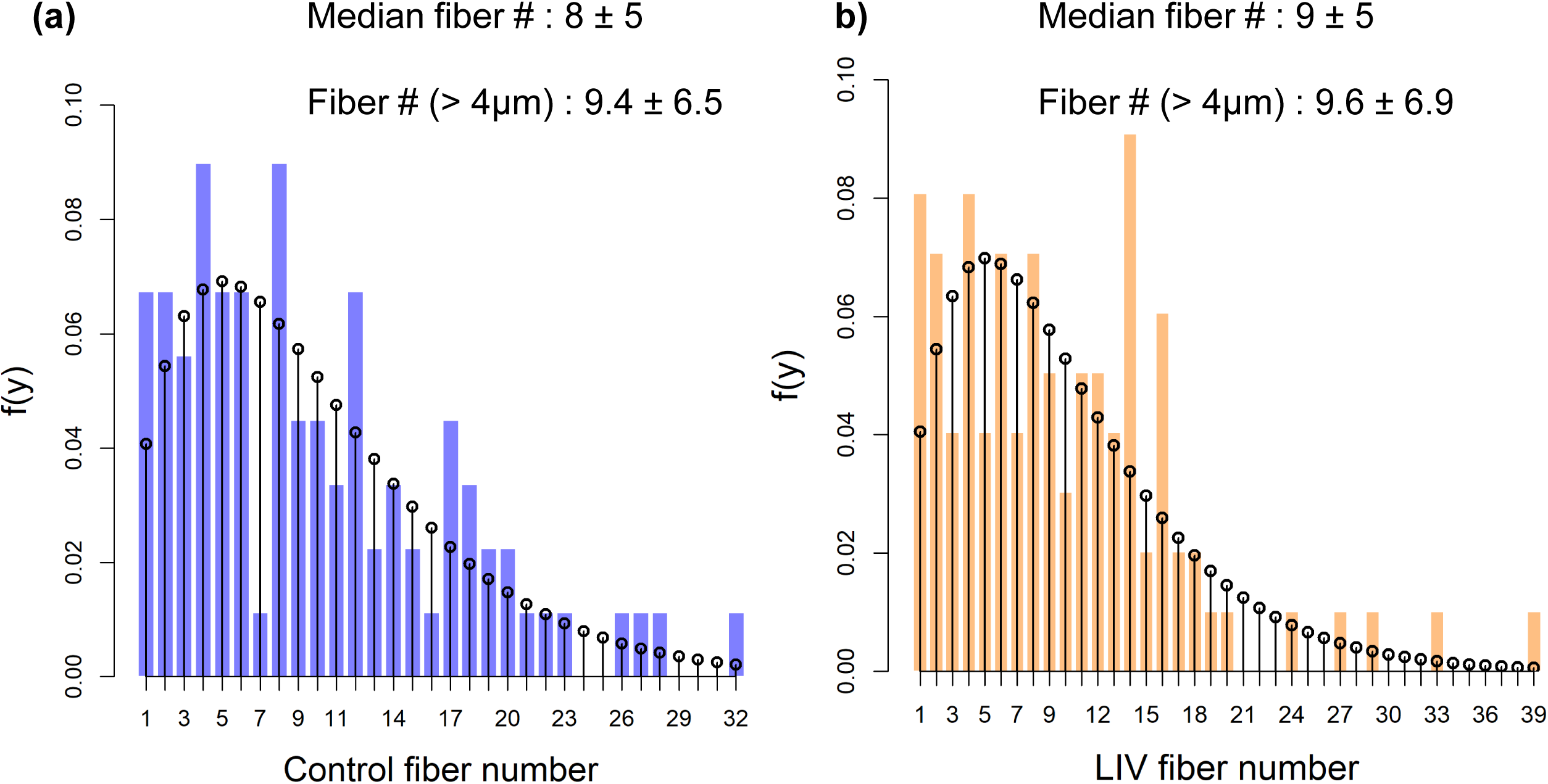
Estimated apical fiber numbers using NBI distribution.

**Figure S7.**
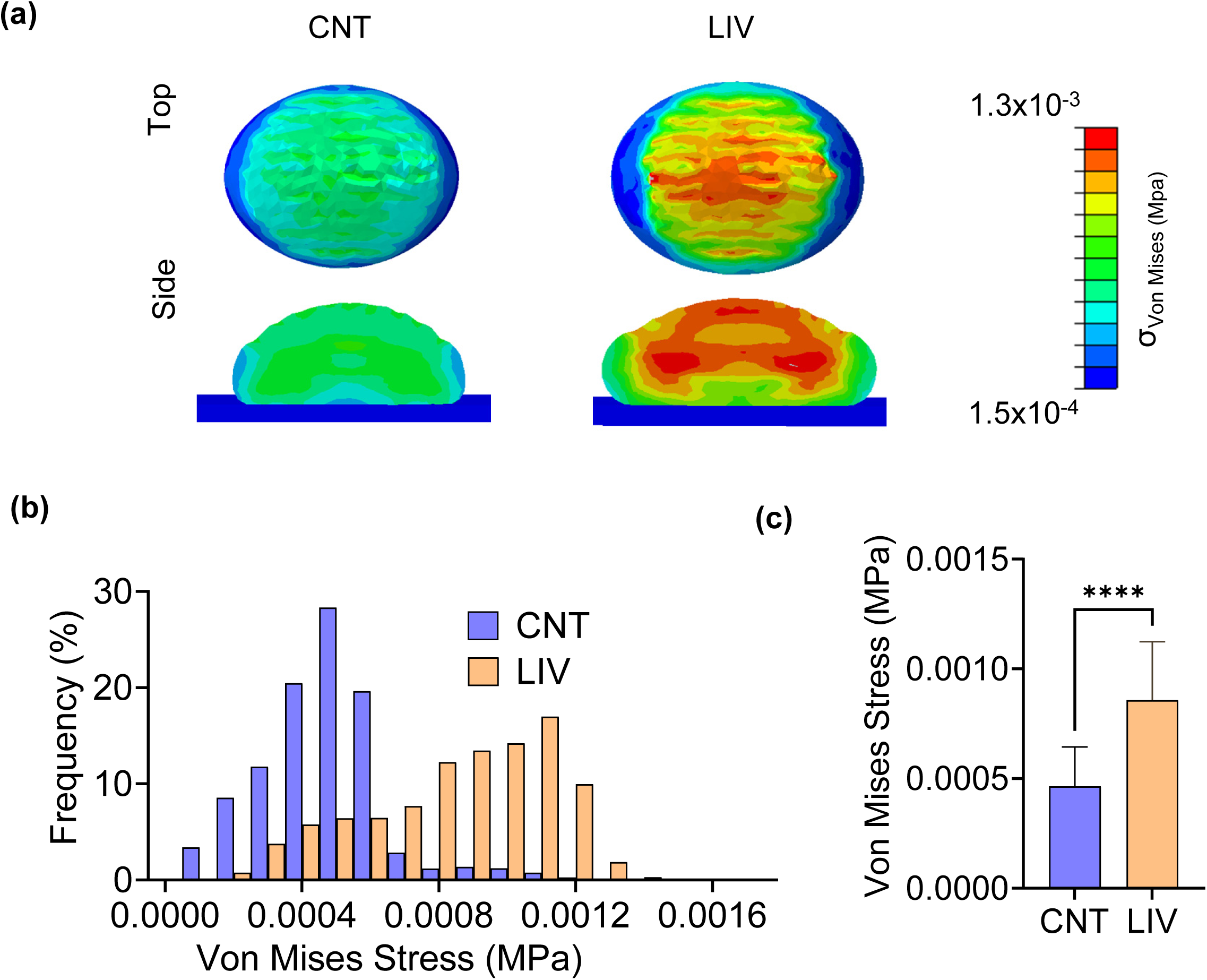
We generated FE models to capture the apical F-actin and nuclear features that best represent both CNT and LIV conditions based on statistical data. We assumed that only longer (>4µm) F-actin fibers that span apical nuclear surface would contribute to the nuclear shape. Utilizing bootstrap confidence interval based on NBI distributions of apical fibers, Using the estimated 9.4 ± 6.5 and 9.6 ± 6.9 apical fibers per cell for CNT and LIV conditions, we implemented a 50% increase in nuclear stiffness as previously reported by our group. Our FE model identified the nuclear surface directly under load bearing actin fibers and mid-nuclei as locations of peak stress. **(b)** With changes to the nuclear stiffness, the FE model predicted that 50% more total F-actin force is required to deform LIV treated nuclei on the apical nuclear surface (135nN) when compared to CNT nuclei (90nN). **(c)** LIV treated nuclei had more elements with higher Von Mises Stress **(d)** increasing the averaged stress by 84% (p< 0.0001) from 0.00043 MPa (0.43kPa) to 0.00096MPa (0.96kPa).

**Supplementary Movie 1:**
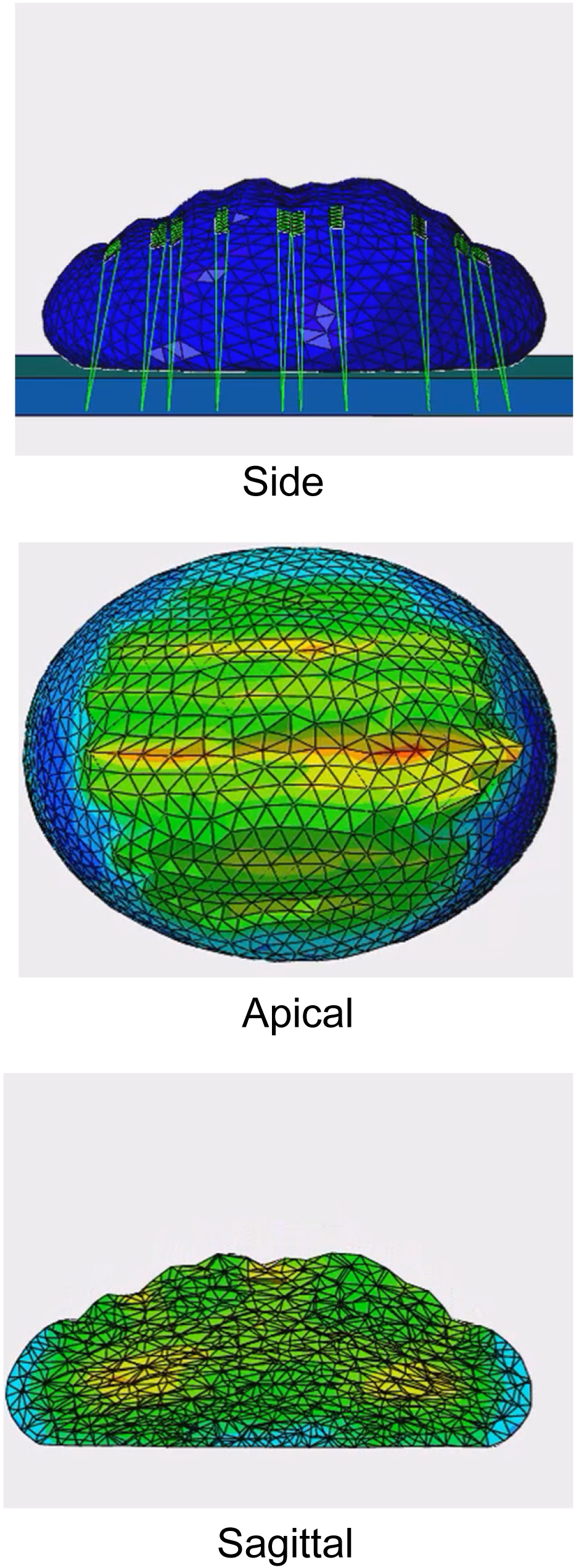
Representative Movie of a FE model run starting from initial conditions to the loaded cell height from side view. Apical and sagittal views also feature Von Mises stresses.

